# Transmigration of *Trypanosoma cruzi* trypomastigotes through 3D cultures resembling a physiological environment

**DOI:** 10.1101/810614

**Authors:** Matías Exequiel Rodríguez, Mariana Rizzi, Lucas D. Caeiro, Yamil E. Masip, Alina Perrone, Daniel O. Sánchez, Jacqueline Búa, Valeria Tekiel

## Abstract

Chaga’ disease, caused by the kinetoplastid parasite *Trypanosoma cruzi*, presents a variety of chronic clinical manifestations whose determinants are still unknown but probably influenced by the host-parasite interplay established during the first stages of the infection, when bloodstream circulating trypomastigotes disseminate to different organs and tissues. After leaving the blood, trypomastigotes must migrate through tissues to invade cells and establish a chronic infection. How this process occurs remains unexplored. Three-dimensional (3D) cultures are physiologically relevant because mimic the microarchitecture of tissues and provide an environment similar to the encountered in natural infections. In this work, we combined the 3D culture technology with host-pathogen interaction, by studying transmigration of trypomastigotes into 3D spheroids. *T. cruzi* strains with similar infection dynamics in 2D monolayer cultures but with different *in vivo* behavior (CL Brener, virulent; SylvioX10 no virulent) presented different infection rates in spheroids (CL Brener ∼40%, SylvioX10 <10%). Confocal microscopy images evidenced that trypomastigotes from CL Brener and other highly virulent strains presented a great ability to transmigrate inside 3D spheroids: as soon as 4 hours post infection parasites were found at 50 µm in depth inside the spheroids. CL Brener trypomastigotes were evenly distributed and systematically observed in the space between cells, suggesting a paracellular route of transmigration to deepen into the spheroids. On the other hand, poor virulent strains presented a weak migratory capacity and remained in the external layers of spheroids (<10µm) with a patch-like distribution pattern. The invasiveness -understood as the ability to transmigrate deep into spheroids- was not a transferable feature between strains, neither by soluble or secreted factors nor by co-cultivation of trypomastigotes from invasive and non-invasive strains. We also studied the transmigration of recent *T. cruzi* isolates from children that were born congenitally infected, which showed a high migrant phenotype while an isolate form an infected mother (that never transmitted the infection to any of her 3 children) was significantly less migratory. Altogether, our results demonstrate that in a 3D microenvironment each strain presents a characteristic migration pattern and distribution of parasites in the spheroids that can be associated to their *in vivo* behavior. Certainly, the findings presented here could not have been studied with traditional 2D monolayer cultures.

**Author Summary:** *Trypanosoma cruzi* is the protozoan parasite that causes Chaga’ disease, also known as American trypanosomiasis. Experimental models of the infection evidence that different strains of the parasite present different virulence in the host, which cannot be always reproduced in 2D monolayer cultures. Three dimensional (3D) cultures can be useful models to study complex host-parasite interactions because they mimic *in vitro* the microarchitecture of tissues and provide an environment similar to the encountered in natural infections. In particular, spheroids are small 3D aggregates of cells that interact with each other and with the extracellular matrix that they secrete resembling the original microenvironment both functionally and structurally. Spheroids have rarely been employed to explore infectious diseases and host-parasite interactions. In this work we studied how bloodstream trypomastigotes transmigrate through 3D spheroids mimicking the picture encountered by parasites in tissues soon after leaving circulation. We showed that the behavior of *T. cruzi* trypomastigotes in 3D cultures reflects their *in vivo* virulence: virulent strains transmigrate deeply into spheroids while non-virulent strains remain in the external layers of spheroids. Besides, this work demonstrates the usefulness of 3D cultures as an accurate *in vitro* model for the study of host-pathogen interactions that could not be addressed with conventional monolayer cultures.

## Introduction

The protozoan parasite *Trypanosoma cruzi* is the etiological agent of Chagas disease, which currently affects about 8 million people. Chagas’ disease is an endemic illness in Latin America that has spread worldwide in the past years. The infection usually develops as a chronic cardiac, digestive or neurologic pathology. The reason why symptoms appear 10 or more years after the initial infection, and only in ∼40% of individuals, remains unsolved, but both host and parasite genetic background should have an impact on the disease outcome. Chagas disease is one of the main health problems in Latin America, causing more than 10000 deaths per year, and incapacity in infected individuals [1].

In humans, the infection initiates with trypomastigotes deposited on mucous or skin, along with triatomine bug faeces, when the insect vector feeds on blood. Trypomastigotes are able to invade any nucleate cells at the infection site. Once inside the cell, trypomastigotes differentiate to amastigotes, which are the intracellular and replicative form. After several division cycles, amastigotes differentiate again into trypomastigotes, the infected cells burst, and parasites are released into the interstitial space. Trypomastigotes can either infect neighboring cells or spread distantly by circulation. Successive cycles of intracellular infection and replication followed by bloodstream trypomastigote dissemination are the hallmark of the initial acute phase, which drives the amplification of the parasitic load, and eventually produces the infection of organs and tissues [2]. The acute phase ends approximately 2-3 months after the initial infection, the time required by the host immune system to control parasitemia and clear most trypomastigotes from peripheral circulation. However, a chronic and persistent infection is already established, characterized by the presence of intracellular amastigotes essentially confined into tissues along with positive serologic tests [2, 3].

The experimental murine model allowed to understand that during the acute phase, trypomastigotes disseminate from the inoculation site to almost all tissues, to render completely parasitized mice, few days after the initial infection [4]. This entails that trypomastigotes should be able to escape from peripheral circulation, cross the vascular endothelium and migrate through the extracellular matrix (ECM) to establish a tisular intracellular infection [5]. The effectiveness of trypomastigotes to cross biological barriers and migrate through tissues will impact on *T. cruzi* ability to produce a severe, moderate or mild tissue infection. This process -that can be linked to parasite virulence and dynamic of infection *in vivo*- is poorly understood. Some authors suggested that the rupture of the endothelial barrier is necessary for the infection of target tissues [6]. On the contrary, others showed that trypomastigotes traverse the endothelial barrier involving a transcellular traffic of trypomastigotes through endothelial cells, mediated by the activation of the bradykinin receptor 2, and without disturbing its integrity nor its permeability [7]. Since different combinations of parasite and mouse strains present differential tissue colonization and target organs of damage [8, 9], the differences between both proposed transmigration models could be attributed to the different strains employed. Either way, trypomastigote transmigration through tissues is an essential event for *T. cruzi* infection. Studies on *T. cruzi* transmigration have been very limited, probably because of the extremely simplicity of monolayers cultures and –on the other hand- the complexity of *in vivo* models, which present low spatio-temporal resolution. Three-dimensional (3D) cultures are physiologically relevant and a good alternative because they mimic the microarchitecture of tissues and can provide an environment similar to the encountered in natural infections [10].

Spheroids are small aggregates of cells that do not adhere to a culture substrate and grow in 3D. Cells interact with each other and secrete the extracellular matrix (ECM) in which they reside, resembling their original microenvironment both functionally and structurally [11, 12]. Spheroids have been broadly employed and deeply contribute to understand mechanisms in cancer biology and immunology [13–15], but they have rarely been employed to explore infectious diseases and host-parasite interactions. Remarkably, the co-culture of spheroids of myocytes with *T. cruzi* trypomastigotes has demonstrated to be an accurate model of fibrosis and hypertrophy that adequately recreates the chronic chagasic cardiomyopathy [16, 17]. In the present work we took advantage of the 3D spheroid technology to disclose how trypomastigotes transmigrate across tissues, which is a key process of the host-parasite interplay in the early steps of infection. We demonstrate that the invasiveness of trypomastigotes from different *T. cruzi* strains and isolates into spheroids can be associated with their *in vivo* behavior and virulence.

## Materials and Methods

### Reagents and sera

Ultrapure Agarose, Carboxy-fluoresceinsuccinimidyl ester (CFSE) and CellTraceTM Far Red (CTFR) were acquired from Invitrogen. Polyethylenimine (PolyAr87-PEI) transfection reagent was obtained from FFyB (University of Buenos Aires). Anti *T. cruzi* antisera developed in mice was generated in our laboratory and used along with goat anti-mouse conjugated to Alexa-488 or Alexa-647 (Molecular Probes).

### Parasites and conditioned media

*T. cruzi* trypomastigotes from different strains and DTUs employed were: SylvioX10, K98, Dm28, 193-733MM and 199-173BB (DTU TcI); Y (DTU TcII); 185-748BB and 186-401BB (DTU TcV) and RA and CL Brener (DTU TcVI). All strains were DNA genotyped by PCR-RFLP of *TcSC5D* and *TcMK* genes (S1 Fig), as described [18].

Parasites were routinely maintained by *in vitro* cultures on Vero cells as previously described [19], and trypomastigotes harvested from supernatants.

Culture-derived trypomastigotes were labeled with CellTrace™ CFSE as we previously described [20]. Trypomastigotes were alternatively labelled with CellTrace™ Far Red CTFR (5 µM), essentially with the same protocol with slight differences in the incubation time (20 min at 37°C, followed by the addition of 1 ml of complete medium and an additional incubation of 5 min at 37°C in the dark). After labelling, the motility of parasites was controlled under light microscope. The percentage of labelled parasites and the fluorescence intensity of CFSE and CTFR was determined by flow cytometry.

*T. cruzi* derived conditioned medium (CM) was obtained by a previously standardized protocol [21]. In brief, cell-derived trypomastigotes (100×10^6^) were washed with PBS, resuspended in 1 ml of MEM without serum (or MEM without parasites as control medium) and incubated for 6 h at 37°C in a 5% CO_2_ humidified atmosphere. Then, parasites were pelleted by centrifugation and the cell-free supernatant (containing both extracellular vesicles as well as vesicle-free secreted material) was centrifuged twice for 10 min at 15000 xg. The clarified supernatant was filtered through a 0.45 μm syringe filter, to obtain the CM, which was aliquoted and stored at −70°C until use.

### Cell culture and stable HeLaR2 cell line generation

Vero, HeLa and HEK293T cells were grown in MEM (Gibco) supplemented with 10% (v/v) fetal bovine serum (Natocor), 100 U/ml penicillin and 10 µg/ml streptomycin. Cells were maintained at 37°C in 5% CO_2_ and 95% air in a humidified incubator. To obtain Hela cells stably expressing LifeAct-RFP, permissive HEK293T cells were first employed to produce lentiviral particles packed with LifeAct-RFP, essentially as described by Gerber *et al* [22]. Briefly, HEK293T were seeded on 24-well plates (3×10^4^ cells/well) and transfected 24 h later by the PEI method with a mix of 0.5 µg pCMV-dR8.9 DVpr (packaging plasmid), 0.05 µg pCMV–VSV-G (envelope plasmid) and 0.5 µg of the pLenti LifeAct-RFP (transfer plasmid) per well. The supernatant containing lentiviral particles packed with LifeAct-RFP was collected at 48 and 72 h after transfection, precleared, and concentrated by centrifugation. HeLa cells (3×10^4^ cells/well) were transduced in MEM containing 10% FBS with a 0.3 m.o.i. of lentiviral particles, and expanded in culture flasks. Cells expressing LifeAct-RFP (HeLaR) were sorted by flow cytometry, and cloned by limiting dilution. The clone number 2 (HeLaR2) stably expressing LifeAct-RFP was selected and employed for all the experiments.

### HeLa spheroids and infection model

Spheroids of HeLaR2 cells were generated by the liquid overlay method [23]. Cells (1000/well) were added to U-bottom 96-well plates coated with agarose 1% in PBS (w/v) and cultured in MEM 10% FBS. The formation of spheroids was controlled by microscopy from 24 h post seeding (S2 Fig). At 72 h, each well contained one spheroid of 300-400 µm diameter, conformed by ≃9000 cells.

For *T. cruzi* infection, twelve spheroids were placed on each well of an agarose pre-coated p24 plate and incubated with 1 or 10 m.o.i. of trypomastigotes (or control medium) for 1 or 24 h, in MEM supplemented with 4% FBS. When appropriate, 100 µl of medium was replaced by 100 µl of CM. When indicated, 2D-monolayers of HeLaR2 cells (10×10^5^ cells/well) were incubated with 1 or 10 m.o.i. of trypomastigotes also labelled with CFSE in a final volume of 500 µl of MEM 4%.

For co-infection assays, 12 spheroids (in one p24 well) were simultaneously incubated with CL Brener and SylvioX10 for 24 h with 10 m.o.i. of each *T. cruzi* strain, labelled with a different stain.

### Cellular infection determined by flow cytometry

Infected spheroids were collected in 1.5 ml tubes, washed three times with PBS and disaggregated by addition of 200 µl 0,25% trypsin/EDTA for 10 min at 37°C. The cellular pellet -collected by centrifugation 10 min at 1000 x g- was washed three times and fixed in PBS 0.5% PFA. Infected 2D-monolayer cells were trypsinized and treated like 3D spheroids. Samples were acquired on a FACSCalibur (Becton Dickinson); gated HeLaR2 by forward and side scatter parameters were selected. A total of 10,000 events were analyzed for each condition. FL1- cells represented uninfected HeLaR2 cells while FL1+ represented cells infected (either with intracellular parasites and/or attached to cell membrane) with CFSE-labelled parasites, FL4+ cells were those infected with CTFR-labelled parasites. Data was analyzed using FlowJo v10.0.7 software. Statistical significance was determined by two-tailed unpaired student *t test* (Prism, GraphPad Software).

### Quantification of free-parasites inside spheroids

Infected spheroids disaggregated by trypsin treatment were centrifuged for 10 min at 5700 x g to collect HeLaR2 cells and parasites that were infecting or attached to HeLa cells, as well as parasites that were free inside spheroids (i.e. not associated to cells but inside spheroids). The pellet was then analysed by confocal microscopy to determine infected cells as well as free parasites (expressed as number of free-parasites for each 100 HeLaR2 cells). Statistical significance was determined by two-tailed unpaired student *t test* (Prism, GraphPad Software).

### Parasitic load into spheroids

The total cargo of parasites inside the spheroids, either infecting cells or free in the ECM, was determined by qPCR. For doing so, each treatment was carried out by duplicate: one sample was used to determine the parasite load associated to spheroids (sample 1) while the other was used to determine the total cargo of parasites in the well (sample 2: parasites associated to spheroids plus parasites free in the medium/well). Infection was carried out as mentioned above. After 24 h, spheroids from sample 1 were collected in 1,5 ml tubes and carefully washed five times with sterile PBS to eliminate parasites from the supernatant avoiding the disassembling of spheroids. Instead, for sample 2 spheroids and medium were collected together, centrifuged for 10 min at 5700 x g and the pellet washed with sterile PBS. Samples were subjected to a standard salting out protocol to obtain genomic DNA [24]. gDNA concentration was measured using Nanodrop and 50 ng were used in each qPCR reaction, which were carried out with Kapa Sybr Fast Universal Kit (Biosystems) in a 7500 Real Time PCR System (Applied Biosystems). The *T. cruzi* single copy gene *PCD6* (TcCLB.507099.50) was amplified with primers v099.50bFw (CAGGCATCACCGTATTTTCCA) and 099.50bRev (CTCTTGTTCCGTGCCAAACA) [25]. To determine *T. cruzi* DNA (*Tc*DNA) abundance, DNA content was normalized to human *GAPDH* gene (53MFZ-GAPDHFw: ACCACCCTGTTGCTGTAGCCAAAT and 54MFZ-Rev: GACCTGACCTGCCGTCTAGAAAAA). Results were analysed with the LinReg software [26]. The percentage of *Tc*DNA inside spheroids was calculated as X%= *Tc*DNAsample1 x 100 / *Tc*DNAsample2 and expressed as mean ± SD of three independent experiments. Statistical significance was determined by student *t test* (GraphPad software).

### Parasite invasiveness and dissemination

Infected spheroids were fixed by adding PFA to a 3.2% final concentration and incubated at 37°C for 1.5 h. Then, spheroids were washed with PBS as described previously [20] and mounted. Fluorescence images were acquired with a confocal Olympus FV1000 microscope. CFSE or CTFR labelled parasites were imaged with a 488 nm or 647 nm laser, respectively, while HeLaR2 cells were imaged at 530 nm. Z-stacks were collected with a 10x objective from 0 to 150 µm in depth with 2 µm intervals in the vertical z-axis. Alternatively, images were acquired with a 40x objective, and the spheroid was scanned at 10, 30 and 50 µm in depth from the surface. To determine the localization of parasites, spheroids were analyzed with a 60x objective and Z-stacks were collected at 0.2 µm intervals in the z-axis. All images were analyzed with ImageJ [27] software; 3D reconstructions and 3D-movies were generated with the 3D-viewer plugin.

### Electron microscopy

Infected spheroids were fixed in 4% PFA and serially dehydrated with increasing ethanol solutions (10-100 %) followed by critical point drying with carbon dioxide. Samples were then coated with 60%/40% palladium/gold and acquired with a scanning electron microscope (Philips - XL Serie 30).

### Free-swimming assay

Trypomastigotes (15×10^6^) were resuspended in 5 ml MEM 4% SFB, transferred to round-bottom centrifuge tubes (Oak Ridge Style) and centrifuged at 2,500 x g for 8 min, which resulted in parasites at the bottom of the tubes forming a thin pellet. The tubes were then incubated 2 h at 37°C, to allow trypomastigotes to freely swim. Aliquots of 1 ml were carefully taken from the top (layer 5) to the bottom; the pellet was resuspended in 1 ml of medium. Parasites in each fraction were enumerated by counting in a Neubauer chamber.

### Statistical analysis

All statistical analyses and graphs were performed with GraphPad Prism 7 (GraphPad Software, USA). We used a two-tailed unpaired *t test* when the means of two groups were compared. When more than 2 groups were compared, we used two-way ANOVA with Bonferroni multiple comparison test. Significant differences were designed when P-value (P) n.s. ≥0.05, *p<0.05, **p<0.01, ***p<0.001.

## Results

### *T. cruzi* trypomastigotes are less infective in 3D spheroids than in 2D monolayer cultures

*Trypanosoma cruzi* presents a high genetic heterogeneity and, currently, *T. cruzi* strains are classified into six clusters or discrete typing units (DTUs), named TcI to TcVI [28]. We selected CL Brener (DTU TcVI) and SylvioX10 (DTU TcI) strains, of high and low virulence, respectively, and whose biologically distinctive behavior in experimental models of *T. cruzi* infection is well characterized [8,29–31].

HeLa cells constitutively expressing LifeAct-RFP (HelaR2 hereon), along with trypomastigotes labelled with CFSE or CTFR were employed to monitor short-time infection dynamics and host-parasite interactions (Fig 1). We first evaluated the infection profile of trypomastigotes both on conventional 2D monolayers and in 3D spheroids (Fig 2). While on conventional 2D monolayer cultures CL Brener and SylvioX10 parasites showed similar infection rates (∼70%) (Fig 2A,C), both strains were much less effective to infect 3D spheroids (Fig 2B,C) and with differences between both strains. Infection with CL Brener rendered higher number of cells with cell-attached or internalized parasites (38,2% CL Brener vs 8,5% SylvioX10), as detected by flow cytometry of disaggregated spheroids (Fig 2B). The total cargo of parasites inside the spheroids, which includes intracellular parasites, surface attached, as well as free parasites migrating inside spheroids through the extracellular matrix, was also higher on CL Brener than SylvioX10 infected spheroids (48% *vs* 18%, determined by qPCR; Fig 2D). Differences between strains were also registered when free-parasites (i.e. not associated to cells) inside spheroids were enumerated (Fig 2E-F). Altogether, these results evidence that both strains have different abilities to infect 3D cultures, being CL Brener strain 2-3 fold more infectious than SylvioX10. These findings were also registered with different multiplicity of infection, and contrast with the similar behavior of both strains on monolayer cultures (S3 Fig).

**Fig 1.**
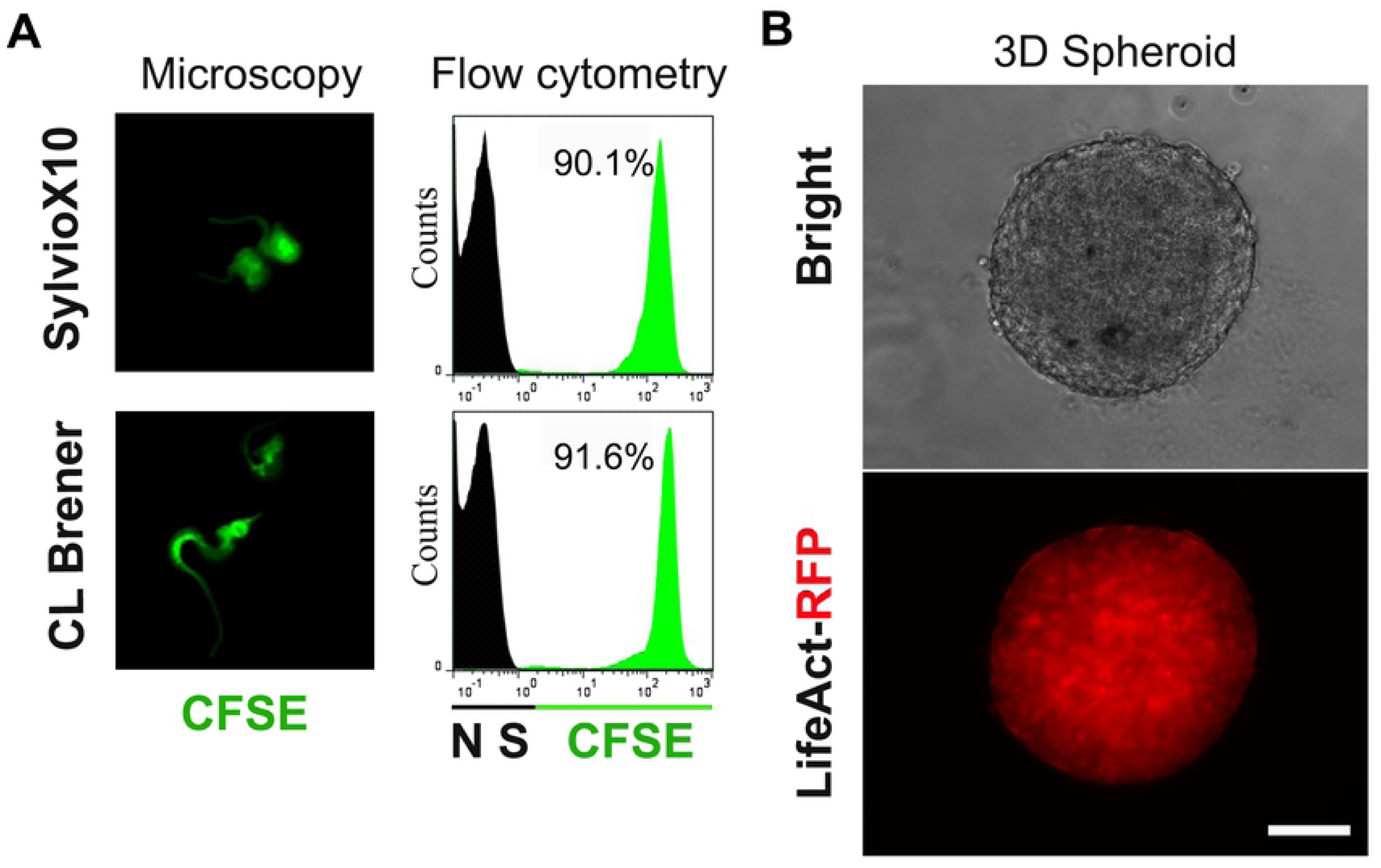
Model setup. (A) *Trypanosoma cruzi* SylvioX10 (DTU TcI) and CL Brener (DTU TcVI) trypomastigotes were labelled with CFSE: fluorescent images (left panel) and quantification by flow cytometry (right panel). Black histograms: non-labelled parasites; green histograms: CFSE labelled parasites. NS: Negative Staining. (B) Spheroid of HeLaR2 cells at 72 h post seeding. Scale bar = 100 µm.

**Fig 2.**
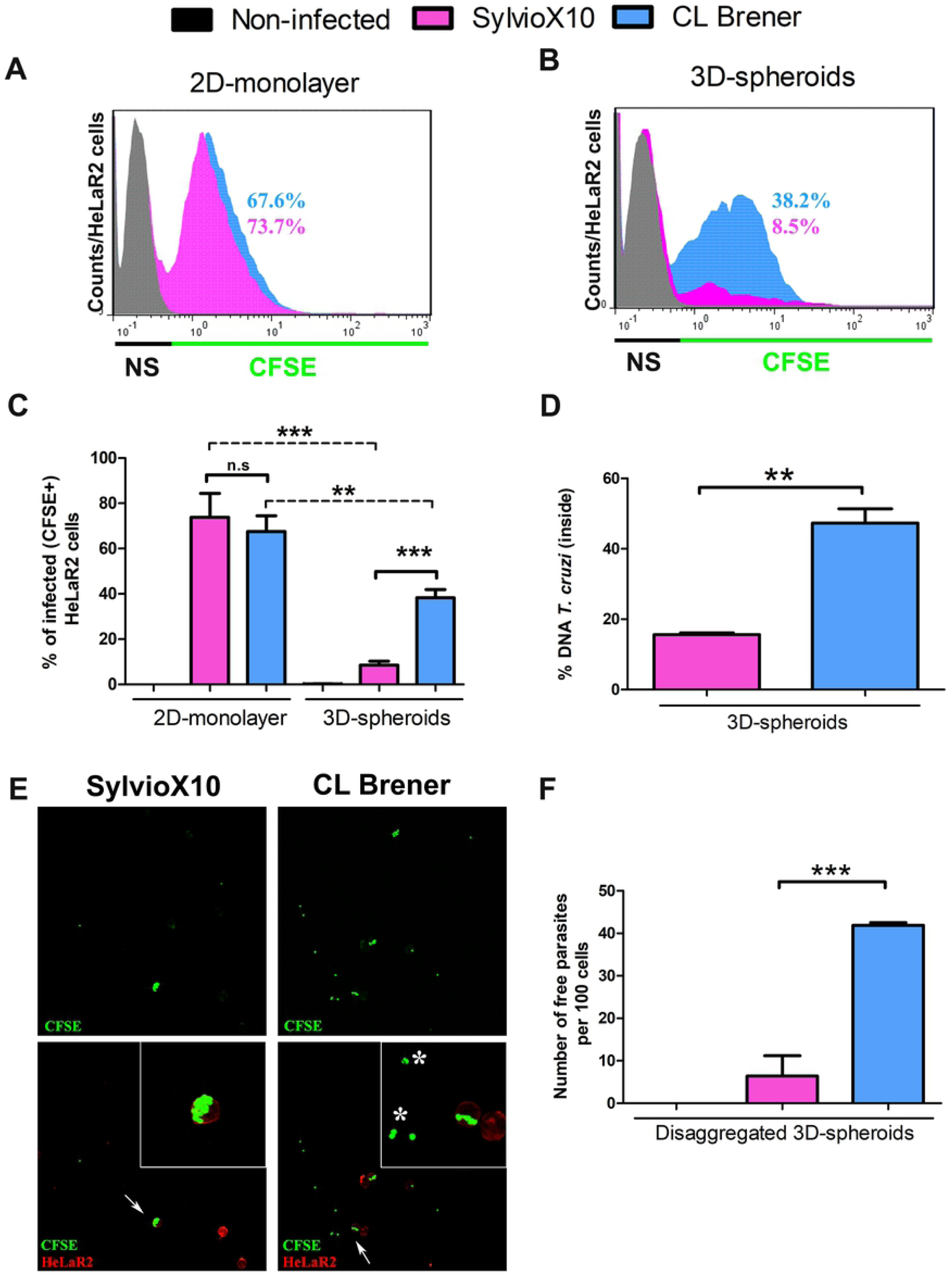
Differential ability of SylvioX10 and CL Brener trypomastigotes to infect 2D-monolayer vs 3D-spheroid cultures. HeLaR2 (grown as monolayers or as spheroids, see Material and Methods section) were incubated with 10 m.o.i. of CFSE-labelled CL Brener or SylvioX10 trypomastigotes or non-infected (control). (A, B) At 24 h post infection, cells were trypsinized and the rate of infected HeLaR2 cells (either with internalized or attached parasites) was determined by flow cytometry. (C) Quantification of three independent experiments carried out as described for A and B. (D) Quantification of the total parasite cargo inside spheroids, which includes both cell-associated parasites and free parasites inside spheroids from three independent experiments. At 24 h post infection either intact spheroids or the whole content of the well (intact spheroids plus culture media with non-internalized parasites) were harvested and total DNA purified. Parasite content was estimated on the basis of qPCR of a single copy *T. cruzi* gene (*PCD6*, TriTryp gene ID: TcCLB.507099.50). Percentage of *T. cruzi* DNA inside spheroids respect to total *T. cruz*i DNA in the well was calculated. (E) Free parasites inside spheroids infected with CL Brener or SylviX10, at 24 h p.i. Representative confocal images of disaggregated spheroids, white arrows: magnified cells; white asterisks: free parasites. (F) Quantification of free parasites released from disaggregated spheroids. Data expressed as number of free-parasites for each 100 HeLaR2 cells. Graphs represent the mean ± SD of three independent experiments *t test,* *p<0.05, **p<0.01, ***p<0.001.

### Trypomastigotes from different strains disseminate differentially inside spheroids

A panoramic view of spheroids, reconstructed from confocal stacks shows that SylvioX10 trypomastigotes were preferentially localized at the spheroid surface. Parasites were mostly focalized in large clumps that resembled a “patch-like” distribution pattern (Fig 3A, S1 movie). By contrast, CL Brener parasites were evenly distributed all over the surface of spheroids (Fig 3B, S2 movie).

**Fig 3.**
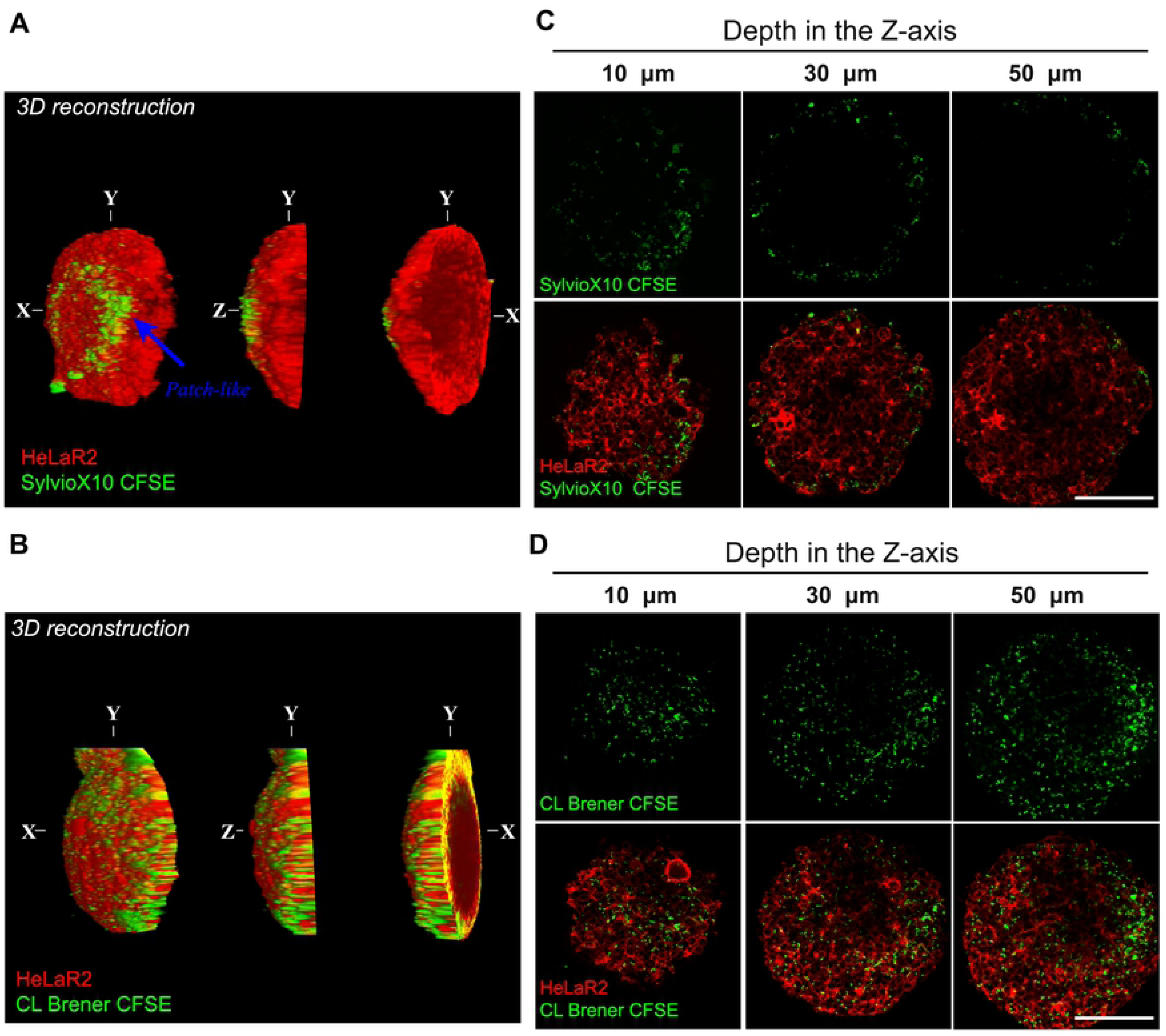
Differential dissemination pattern within spheroids of different strains of *T. cruzi*. (A and B). 3D reconstruction of HeLaR2 spheroids infected with CFSE-SylvioX10 or CFSE-CL Brener trypomastigotes. Z-stack images were obtained by confocal microscopy. The distribution on the surface (X-Y left image), the side plane (Z-Y middle image) and inside the spheroid (Y-X right image, transversal view) are shown. Green: CFSE-trypomastigotes. Red: LifeAct-RFP of HeLa cells. (C and D) Representative images of three confocal planes obtained at 10, 30 and 50 µm in depth on the z axis (40x objective). The detailed distribution pattern of parasites is observed -green fluorescence-. Scale bar: 100 µm.

The transmigration and invasiveness of trypomastigotes was analyzed by scanning the spheroids by confocal microscopy. Most SylvioX10 trypomastigotes were retained at spheroid surface or at the first layers of cells, and only scarce trypomastigotes were detected up to 30 μm in depth (Fig 3C). On the other hand, CL Brener parasites were able to deepen into spheroids: migrated uniformly and were easily detected up to 50 μm in depth (Fig 3D). The migration through the spheroid seems to be a fast movement because similar patterns were observed from 1 h post infection (S4 Fig). In brief, confocal scanning evidenced that CL Brener trypomastigotes can efficiently transmigrate deeply into spheroids, while SylvioX10 is retained at the surface, which corresponds with the differential virulence of both strains.

### Trypomastigotes of high migratory CL Brener strain use a paracellular migration route to move inside spheroids

To answer how trypomastigotes spread within spheroids, we analyzed the parasite-spheroid interaction with higher resolution techniques, such as scanning electron microscopy (SEM) and higher power snapshots by confocal microscopy.

As evidenced by confocal microscopy, SEM images also showed numerous SylvioX10 parasites attached to the surface of individual cells (Fig 4A, panels b, c; white arrows). Notably, CL Brener trypomastigotes were predominantly caught entering into spheroids through the space between cell-cell junctions (Fig 4A, panels e, f; white asterisks). Spheroids infected with SylvioX10 also presented multiple intracellular amastigotes in the superficial layers -first 10 µm- of cells with an untidy distribution (Fig 4B panels a, b and c; 4C and S3 movie). Confocal slices of spheroids infected with CL Brener made evident that there is an orderly distribution pattern around cell-cell contacts (Fig 4B panels d, e, f; white asterisks). At shorter times, CL Brener trypomastigotes were plainly observed in the space between cells, on their way through the extracellular matrix, suggesting a paracellular route of transmigration within the spheroid (Fig 4D and S4 and S5 movies). CL Brener trypomastigotes were also detected intracellularly, though fastened to the cellular membrane (S6 movie).

**Fig 4.**
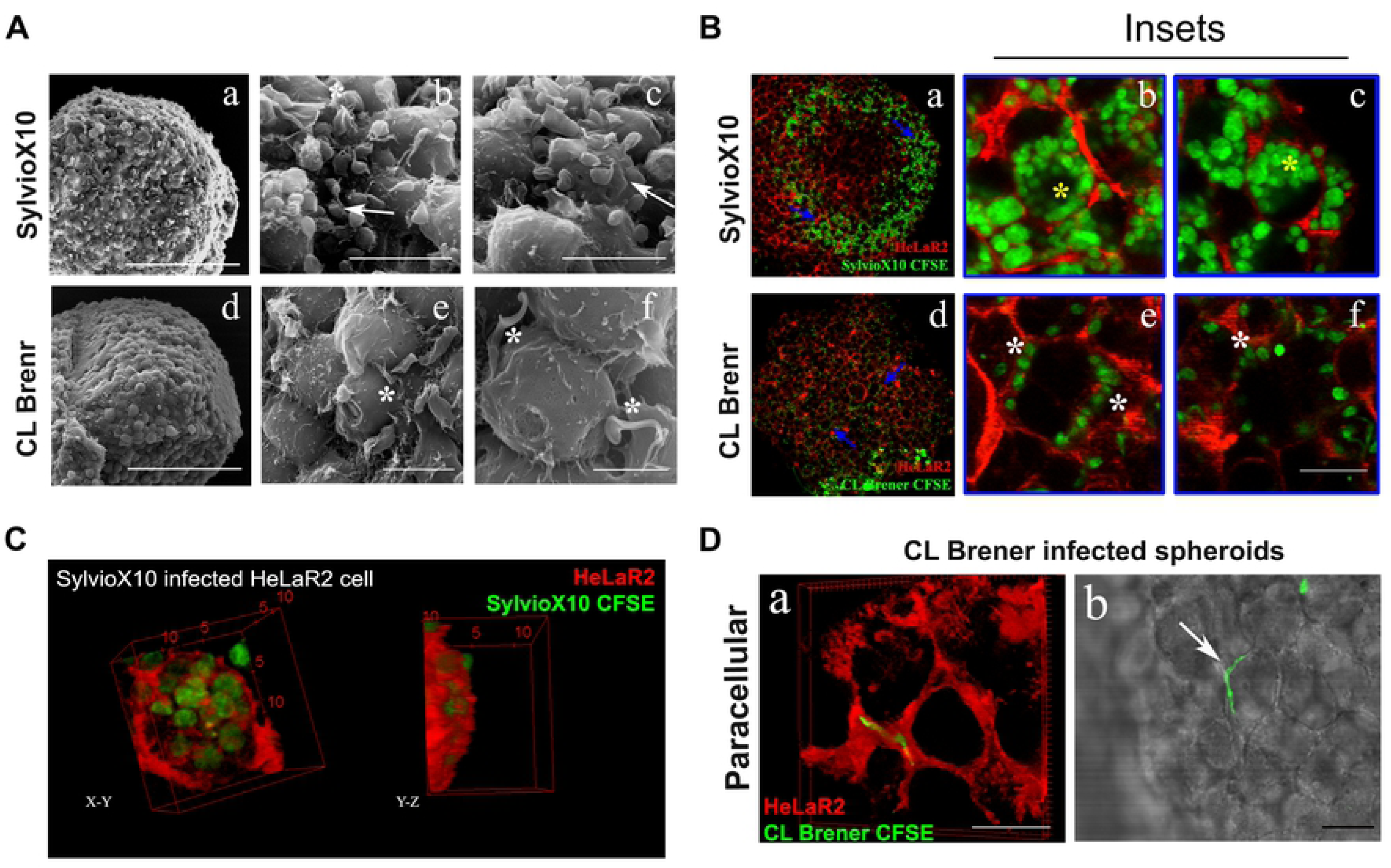
Differential host cell-parasite interaction pattern of different strains of *T. cruzi* on the spheroid surface. Spheroids were cultured with CFSE-SylvioX10 or CFSE-CL Brener. (A) Cell-parasite interaction on the surface of infected spheroids after 24 h of infection. SEM microscopy showing the whole surface of infected spheroids (images a and d) or detailed cell-parasite interactions (b-c for SylvioX10 and e-f for CL Brener) are shown. White arrows show groups of parasites on the surface of SylvioX10 infected spheroids. White asterisks show CL Brener parasites located in the site of cell cell-cell contact. Scale bar for a and d: 100 µm; b and c: 10 µm; d: 100 µm; e: 10 µm: f: 5 µm. (B) Confocal microscopy capturing cell infection at 10 µm of infected spheroids with 60x objective. The distribution pattern of parasites on the surface of infected spheroids is observed in a and d images for SylvioX10 and CL Brener, respectively. Blue arrows show magnified insets (b-c for SylvioX10 and e-f for CL Brener). Yellow asterisks show multi-infected cells. White asterisks show cell-cell contact associated parasites. Scale bar = 15 µm. (C) Multi-infected cell on the surface of SylvioX10 infected spheroids was reconstructed in 3D. Multiple intracellular amastigotes can be observed. (D) Spheroids of HeLaR2 cells were incubated with CFSE-CL Brener trypomastigotes for 1h and then photographed by confocal microscopy. CFSE-trypomastigotes in the paracellular space both in fluorescence images (a) as well as in bright light (b) are shown. Scale bar 15 µm.

### The capacity of transmigration is not transferable between strains

CL Brener and SylvioX10 strains presented not only dissimilar invasiveness profiles, but also their allocation at the superficial layers of spheroids was very distinctive (Fig 3 and 4). We then investigated if transmigration could be transferred from the highly migrant CL Brener strain to the low migrant SylvioX10, through soluble or secreted factors or by co-cultivation of both strains (Fig 5). Interestingly, each strain retained its own dissemination pattern (Fig 5A) and rate of infection (Fig 5B) irrespective of the presence (or absence) of the other strain. Conditioned media (including both soluble and vesicle-contained secreted factors) from CL Brener or SylvioX10 did not cause changes in the invasiveness pattern or percentage of infected cells (S5 Fig). Together, these results indicate that the transmigration capacity of *T. cruzi* is a strain-specific trait that cannot be transferable by soluble or secreted factors, nor through co-cultivation of migrant and non-migrant trypomastigotes.

**Fig 5.**
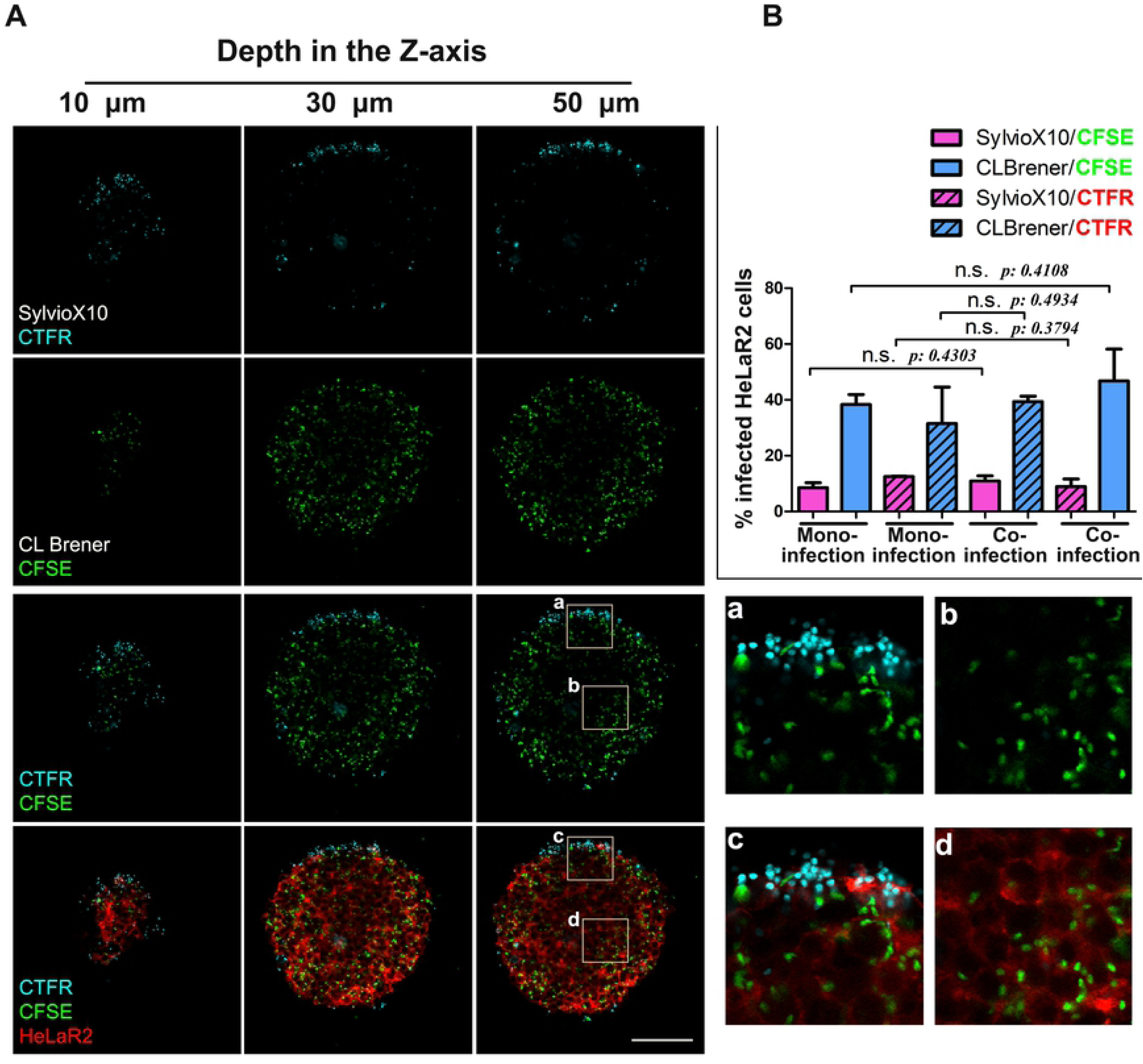
Invasiveness within spheroid is an intrinsic feature of each strain, not complemented in trans. Spheroids of HeLaR2 cells were incubated with CFSE-CL Brener, CTFR-CL Brener, CFSE-SylvioX10 or CTFR-SylvioX10 or simultaneously co-incubated with both strains labelled with different dyes. (A) Representatives images of three confocal planes obtained at 10, 30 and 50 µm in depth -on the z axis-with a 40x objective. Cyan: CTFR-SylvioX10; green: CFSE-CL Brener; red: HeLaR2 cells. Magnified insets (a-d) are shown in the right panels. Scale bar = 100 µm. (B) The same assay as described in A but spheroids were disaggregated and cells with either attached or internalized parasites were quantified by flow cytometry. Infections with one (mono-infections) or both strains simultaneously (co-infections) were carried out also with interchanged dyes. The percentage of infected cells was significantly different between SylvioX10 and CL Brener parasites in all conditions tested. On the other hand, no differences in the infection rate for each strain, both with the use of different staining dyes, as well as, during the mono infections versus the co-infections were observed. Graphs represent the mean ± SD of three independent experiments. Data were analyzed by ane-way ANOVA followed by Bonferroni’s multiple comparison test.

### Transmigration deep inside spheroids can be linked to virulence

To evaluate if the differential transmigration profile could be linked to parasite DTU or virulence, the transmigration into spheroids of other well characterized *T. cruzi* strains was also analyzed. Low virulent strains (K98 and Dm28; TcI) presented low ability to transmigrate into spheroids (Fig 6). In contrast, virulent strains (RA [TcVI] and Y [TcII]) showed a transmigration pattern resembling the observed for CL Brener (Fig 6) (Fig 6A and S7 and 8 movies).

**Fig 6.**
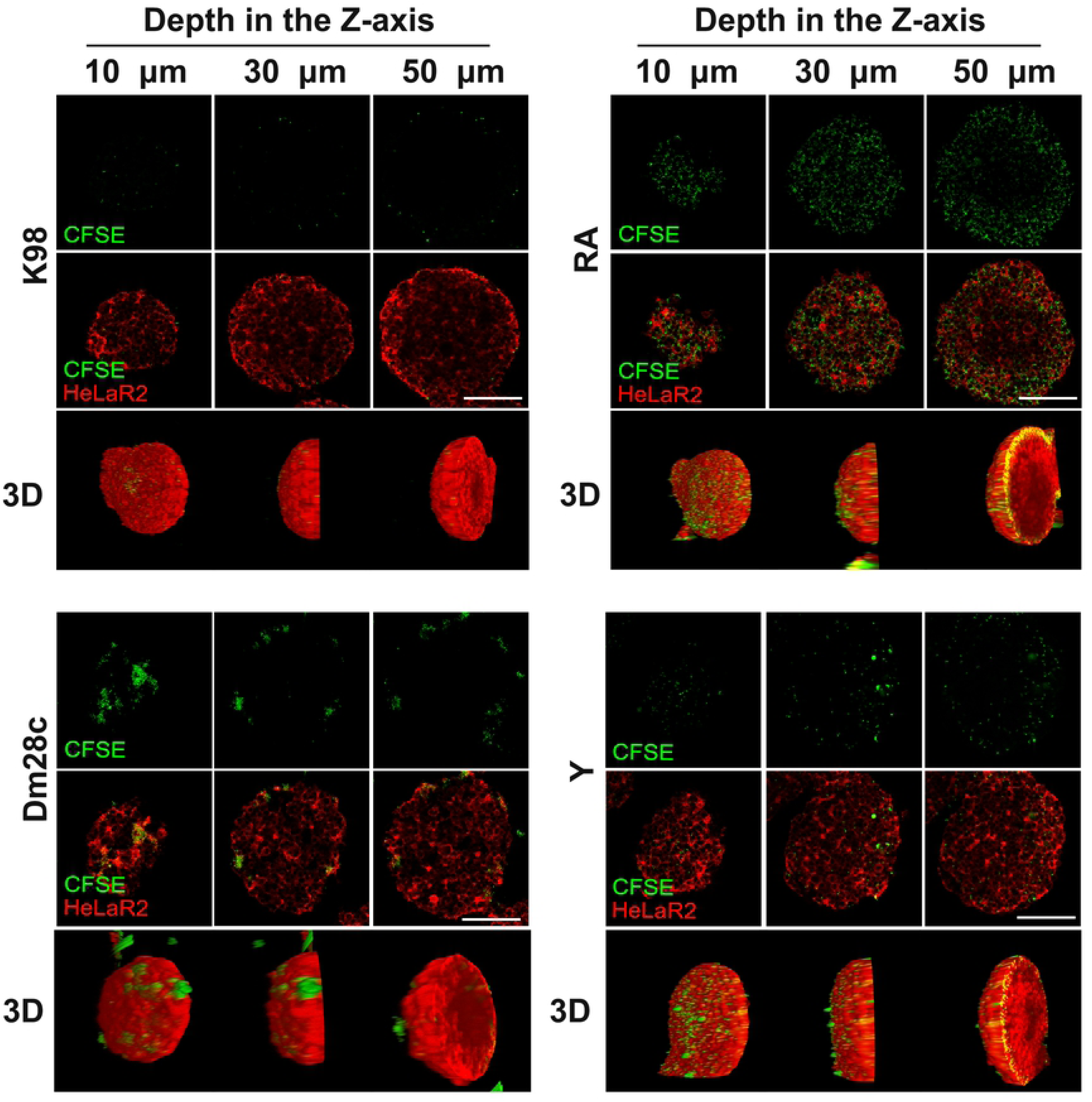
The capacity of dissemination is linked to virulence. Spheroids of HeLaR2 cells were incubated with k98, Dm28c, Y or RA strains for 24 h. Three confocal planes were obtained at 10, 30 and 50 µm in depth -on the z axis-with 40x objective. The detailed distribution pattern of parasites is observed. 3D-reconstruction is shown in lower panels. Green: CFSE labeled parasites. Red: LifeAct-RFP. Scale bar = 100 µm.

Finally, we analyzed the transmigration of four recent clinical isolates of *T. cruzi*. One isolate was derived from a *T. cruzi*-infected mother that after several pregnancies never delivered an infected child (isolate 773MM, TcI; non-congenital transmission). The other isolates, derived from babies that were born congenitally infected (isolates 173BB, TcI; 748BB, TcV; and 401BB, TcV; congenital transmission) [32]. The non-congenital isolate (733MM) [32] showed a low ability to migrate deep inside spheroids. Also, a “patch-like’ distribution pattern, similar to the observed with SylvioX10 strain was observed (Fig 7B). In contrast, congenitally isolated parasites presented a highly migrant phenotype. Either DTU TcI or TcV isolates from congenitally infected babies were found deeply inside spheroids and easily visualized along the first 50 µm in depth (Fig 7A).

**Fig 7.**
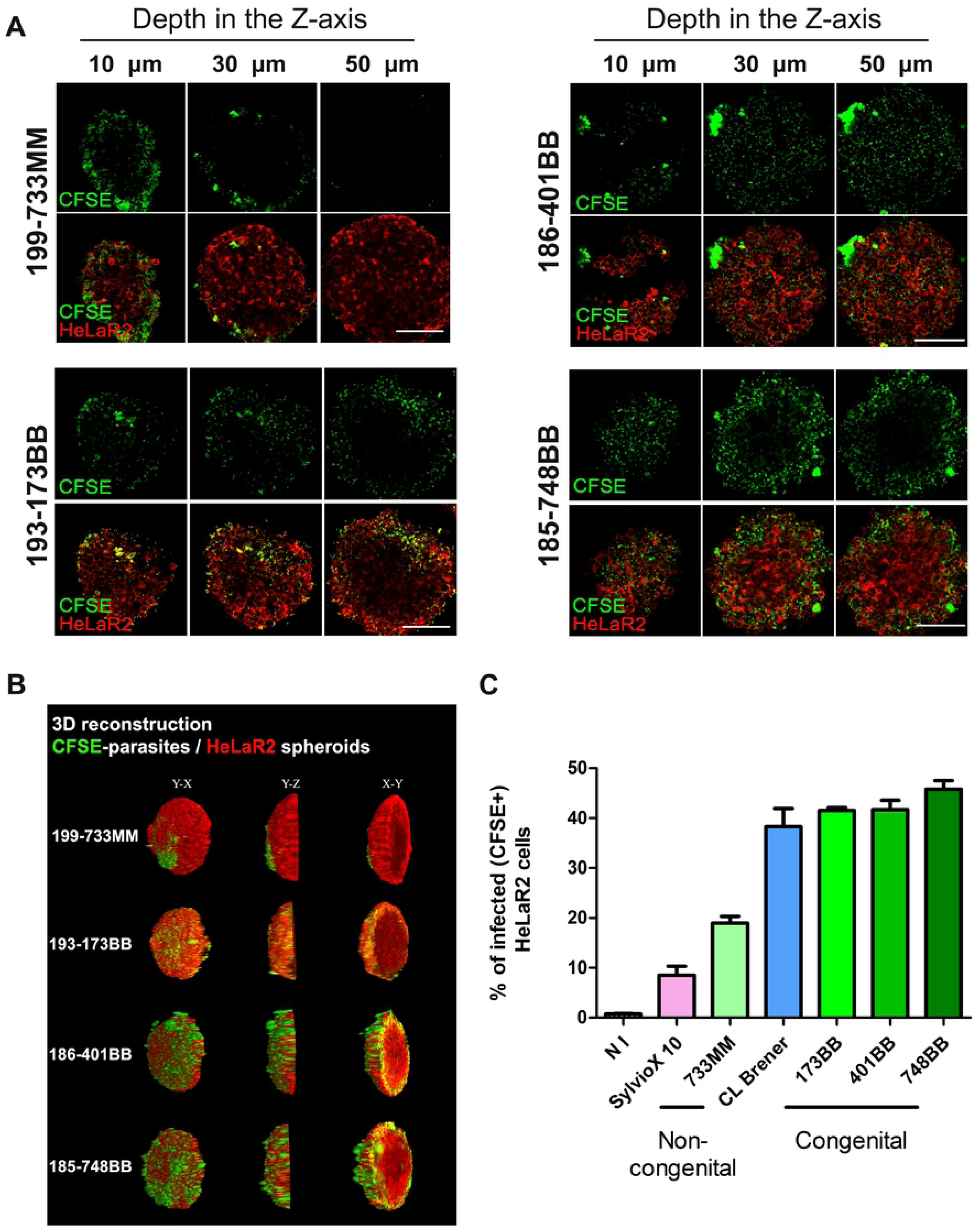
Congenital trypomastigotes are more invasive and infective than non-congenital parasites. Spheroids of HeLaR2 cells were incubated with 733MM, 173BB, 401BB or 748BB recent clinical isolates for 24 h. (A) Confocal planes were obtained at 10, 30 and 50 µm in depth -on the z axis-with 40x objective. The detailed distribution pattern of parasites is observed. Sacle bar = 100 µm. (B) Three-D-reconstruction of infected spheroids with each strain is shown. Green: CFSE labeled parasites. Red: LifeAct-RFP. (C) Percentage of infected HeLaR2 cells to compare infectivity of non-congenial vs congenital isolated. Pink and blue bars show SylvioX10 and CL Brener % of infection, respectively, as comparison points. Graphs represent the mean ± SD of three independent experiments.

The cellular infection produced by 733MM was ∼20%, a value near the one registered with the low virulent SylvioX10 strain, while the 40% of infection of cells in spheroids produced by congenital isolates resembled the infection produced by CL Brener strain (Fig 7C).

Ultimately, because trypomastigotes of different strains behave differently when they are allowed to swim freely in the medium, we analyzed parasite motility as a possible trait linked to invasiveness within spheroids (S6 Fig). We consider a strain (or isolate) as poor motile when more than 20% of the parasites remain at the pellet in this assay. Although the swimming ability of SylvioX10 and CL Brener strains were considerably different and agree with their behavior in spheroids, other strains showed no association between the transmigration inside spheroids and their swimming ability (for example, up to 60% of parasites from 173BB and 401BB -congenital isolates highly migrant in spheroids-remained between the pellet and layer 1).

## Discussion

The infection with the protozoan parasite *Trypanosoma cruzi* evolves from a short acute to a long lasting chronic phase when cardiac, neurological or intestinal disorders become evident [3]. Although the pathology appears only at the chronic phase, the infection of tissues initiates during acute phase and is the consequence of the early dissemination of trypomastigotes. Indeed, *T. cruzi* can disseminate and establish an intracellular infection in any tissue of the mammalian host [5]. To accomplish this, trypomastigotes must migrate and actively cross several biological barriers, from the initial infection site to the target organs of damage, where parasites replicate intracellularly as amastigotes [2]. Murine experimental models of *T. cruzi* infection have helped to understand that some parasite strains present tropism for certain tissues or organs while others are essentially pantropic and can colonize indistinctly any tissue [5,33,34].

Usually, pantropic strains are more virulent in the murine model. It can be inferred that, since those strains colonize a broader range of tissues, they are also more efficient in the transmigration process. However, how trypomastigotes transmigrate, the mechanisms underlying this process and its significance in the host-parasite interplay are poorly understood.

In this work, we employed 3D cultures to mimic the tisular microarchitecture encountered by trypomastigotes in the mammalian host during its *in vivo* life cycle. We studied the process of transmigration and dissemination of the parasites across spheroids for the first time, and demonstrate a link between 3D transmigration and *in vivo* behavior. Strains or isolates that are more virulent *in vivo* (in natural or experimental infections) transmigrated deeper inside spheroids than no virulent strains. In an *in vivo* infection, the ability to transmigrate will favour pathogen dissemination into the host, at the same time that parasites evade the immune system and increase the opportunity to find an adequate microenvironment to settle for the tissular infection [35–38].

By employing the 3D spheroid model, we focused on one hand in the ability of *T. cruzi* strains to infect mammalian cells (evaluated by flow cytometry as cells with either internalized or attached parasites). On the other hand, we also examined the invasiveness of trypomastigotes, which means how deep inside the spheroids trypomastigotes are detected. Somehow both events (invasiveness and infection) are linked by the fact that *T. cruzi* strains that were highly migrant were also those that presented higher infection rates, probably because the transmigration was a necessary step to infect the cells located deep inside the spheroids. This fact can also explain why poorly migrant strains presented low infection rates in the 3D model, irrespective of their accurate infection rate in conventional 2D monolayer cultures. However, considering the times at which transmigration was analyzed, it is unlikely that transmigration was the result of cellular invasion and replication of parasites. We postulate the transcellular and paracellular transmigration routes as two possible ways for trypomastigotes to reach the deeper layers of spheroids. Even more, we speculate that how *T. cruzi* transmigrates can be also a strain dependent trait and that different strains or isolates can employ differential transmigration strategies. Electronic microscopy images strongly suggest that CL Brener strain goes through spheroids by a paracellular route, without crossing the cells but between cell-cell junctions. Although the biological significance of this transmigration strategy should be carefully studied, we hypothesize that the paracellular route would allow the parasite to internalize into the tissues without disrupting the cellular homeostasis and, therefore, without triggering an inflammatory response. Moreover, a paracellular route would be a faster transmigration mechanism for trypomastigotes to find their target allocation inside tissues, without the need to invade and replicate intracellularly. In line with these results, Coates et al (2013) showed that *T. cruzi* trypomastigotes can cross a monolayer of endothelial cells without cell damage. They suggested that this process might be mediated by the protease cruzipain, which can convert kininogen to bradykinin (involved in endothelial permeability) [7]. The picture of trypomastigotes distribution was very distinctive between CL Brener and SylvioX10 strains, even from the initial steps of interaction with mammalian cells in our 3D model. While CL Brener trypomastigotes are regularly distributed and positioned in between cell-cell junctions on the external layers of the spheroid surface, SylvioX10 trypomastigotes are grouped in patches of several parasites stuck over the cells. Previous works with *Trypanosoma brucei* evidenced that trypomastigotes can cross the blood brain barrier both by transcellular and paracellular routes, promoting the expression of ICAM-1 and VCAM-1 [39–41]. On the other hand, *T. gondii* employs a paracellular route for tissue transmigration, through the interaction between TgMIC2 and host occludins from TJs and ICAM-1[42, 43]. Interference with the transmigration process avoids *in vivo* infectivity both in *T. gondii* and *P. falciparum* [44, 45]. Interestingly, the loss of genes associated with transmigration in *P. falciparum* did not impair cellular invasion, supporting the idea that tissue invasiveness and cellular infection can be two independent processes [45].

Trypomastigote’s motility can be understood as its ability to present a directional movement, which in turn could impact on the cellular infection rates. We found a pronounced difference in swimming motility between SylvioX10 (low migrant and low motile) and CL Brener (high migrant and high motile) trypomastigotes. However, analyses of a broader panel of strains demonstrated that transmigration cannot be solely explained by the motile ability of trypomastigotes. Presumably, transmigration depends both on motility and migration of the parasite as well as on its interaction with the surrounding microenvironment. In this sense, Barragan *et al*, (2002) showed that a high migration rate of *T. gondii* is associated with a highly virulent phenotype [46]. Indeed, correlation between high virulent strains and congenital toxoplasmosis has also been noted [47, 48]. On the other hand, Éva Dóró et al (2019) have very recently evaluated the migration of trypomastigotes of *T. carassii* in an *in vivo* model. This awesome work clearly shows that the movement of parasites inside zebrafish occurs through the interstitial space and how its density and compaction determines the direction of trypomastigotes migration [49].

During congenital transmission of *T. cruzi* infection, trypomastigotes must cross epithelial and connective tissues that compose the placental barrier to gain access to and infect the fetus. Therefore, transplacental infection is another aspect of *T. cruzi-*host interplay that is associated with parasite transmigration. We have characterized the invasiveness inside spheroids of isolates recently obtained from babies born with congenital Chagas disease. We demonstrated that the congenital isolates were highly invasive into spheroids, in contrast with the isolate obtained from a mother, which after delivered several children never transmitted the infection to her offspring, which showed a low/moderate transmigration ability. *T. cruzi* congenital transmission is the result of a complex interaction between trypomastigotes and the placental barrier [50–52]. Recently, Juiz et al (2017) described a differential placental gene response induced by strains with different tropism and virulence. They reported that a strain that was isolated from a human case of congenital infection (VD) presented higher tropism by the murine placenta than a non-virulent and myotropic strain (K-98) [53]. In our 3D model, K-98 strain showed a low migrant phenotype. Although we did not analyze the transmigration profile of VD strain, all the congenital isolates assayed here were highly migrant.

The intratisular migration is key during the development of metastasis and it has been approached in several studies on cancer [54]. Tumoral cells produce and prompt to the secretion of cytokines and proteases that will favour the migration across different biological barriers. Proteases are necessary to disrupt cell-cell junctions, extracellular matrix and the basal lamina [55]. Although little is known about the transmigration process in host-pathogen interactions, the secretion of proteases could also be required to disrupt intercellular junctions and extracellular matrix for *T. cruzi* transmigration. However, in our experimental conditions, the invasiveness was not a transferable feature between strains, neither by soluble or secreted factors nor by co-cultivation of invasive and non-invasive trypomastigotes. This observation suggests that unsecreted and strain specific factors are required to transmigrate into the spheroids, while it does not exclude the involvement of proteases and other soluble factors. Differentially-expressed and/or strain specific membrane-associated molecules from trypomastigotes might be targets to be evaluated in the near future.

Altogether, our results demonstrated that in a 3D microenvironment each strain presents a characteristic migration pattern and tissular distribution that could be associated to their *in vivo* behavior. Our work also validates the accuracy and utility of the 3D spheroid model to study complex host-parasite interactions. Certainly, the findings presented here could not have been studied with traditional 2D monolayer cultures.

## Acknowledgements

We acknowledge L. Sferco and A. Chidichimo for technical assistance with *in vitro* parasite cultures, and V. Campo and G.V. Levy for advice in qPCR assays. We are grateful to A.M.M. Massaldi for checking the English version of the manuscript. This work was supported by grants from Agencia Nacional de Promoción Científica y Tecnológica / Fondo para la Investigación Científica y Tecnológica -ANPCyT/FONCyT- (PICT-2016-0108 to VT and PICT 2017-2644 to MER) from Argentina. The funders had no role in study design, data collection and analysis, decision to publish, or preparation of the manuscript.

## Supporting information captions

**S1 Fig. Genotyping of *T. cruzi* strains and isolates** (according to Cossentino et al, 2012).

(A) TcSC5D amplification product was digested with SphI/HpaI restriction enzymes and fragments resolved in 2% agarose gels. Fragment length polymorphism defined lineages TcI for SylvioX10, K98, 733MM and 173BB; TcII for Y; TcV/VI for CL Brener, RA, 401BB and 748BB. (B) The digestion of TcMK product with XhoI allows to distinguish between DTUs TcV (401BB and 748BB) and TcVI (CL Brener and RA).

**S2 Fig. The 3D culture model.**

(A) Monolayer of HeLa cells stably expressing LifeAct-RFP (HeLaR2), generated by lentiviral infection of the parental HeLa cells. (B) Spheroids formation by liquid overlay. After 24 h, spheroids are irregular aggregates of cells. From 72 h after culturing, spheroids seem to be compact aggregate of cells with well delimited borders.

**S3 Fig. Infection of 2D-monolayer and spheroids at m.o.i. 1, with strains CL Brener and Sylvio.**

HeLaR2 cells were incubated with CFSE-labelled trypomastigotes for 24 h and then monolayers or spheroid cultures fixed. (A) the Infected cells were determined by flow cytometry, which detects both attached and internalized parasites. (B) Quantification of 3 independent experiments, carried out as described for A. (C, D) Spheroids were disaggregated and released parasites (free parasites inside spheroids) were visualized by fluorescence microscopy and quantified. * *t test* <0.05.

**S4 Fig. Dissemination pattern within spheroids of CL Brener after 1 h of infection.**

Spheroids incubated with 10 m.o.i. of CFSE-CL Brener parasites for 1h were fixed and confocal planes were obtained at 10, 30 and 50 µm in depth -on the z axis- with 40x objective. Representative images of the invasiveness of CL Brener parasites -green fluorescence- that were detected even at 50 µm in depth within the spheroids. Scale bar = 100 µm.

**S5 Fig. Invasiveness within spheroid is not mediated by soluble factors.**

(A) spheroids were incubated with CFSE-CL Brener (left) or CFSE-SylvioX10 (right) trypomastigotes for 24 h along with control media, or conditioned media (CM) from either SylvioX10 or CL Brener trypomastigotes. Then, spheroids were fixed and confocal images were taken at 10, 30 and 50 µm in depth from spheroid surface. (B) Spheroids incubated with CL Brener or SylvioX10 trypomastigotes in presence of control media or conditioned media were disaggregated with trypsin and % of infected cells was measured by flow cytometry.

**S6 Fig. The free motility *in vitro* is not associated with invasion inside spheroids.** Trypomastigotes were pelleted in round bottom tubes and then incubated for 2 h at 37°C in medium to allow free-swimming (Left image). Aliquots of 1 ml (layer 1 to 5) of the medium from top to bottom were carefully collected and quantification of the % of parasites in each layer and pellet was carried out by microscopy, on Neubauer chamber (graph). Isolates / strains are classified in two groups: those that presented low motility [>40 % of trypomastigotes in the pellet after 2 h] and those with high motility [<20% of parasites in the pellet] and trypomastigotes were homogeneously distributed among all layers]. As a reference, the low migran strains (into spheroids) are in pink and the high migrants are in sky blue. pasa a suplementaria

**S1 and S2 movies.**

HeLaR2 spheroid infected with CFSE-SylvioX10 (video1) or CFSE-CL Brener (video 2); 3D reconstruction. Z-stack images were obtained by confocal microscopy using 10x objective. The videos were realized by using ImageJ software -3D viewer plugin-. Half of an infected spheroid rotating on the Y axis is shown. Green: CFSE-parasites. Red: LifeAct-RFP of HeLa cells.

**S3 movie.**

3D reconstruction of a single HeLaR2 cell on the surface of a SylvioX10-infected spheroid. Z-stack images were obtained by confocal microscopy using 60x objective. The video was realized by using ImageJ software -3D viewer plugin-. Multiple amastigotes within a single cell are shown. Green: CFSE-parasites. Red: LifeAct-RFP of HeLa cells.

**S4 movie.**

3D reconstruction of a single CL Brener trypomastigote located inside the spheroid (≈30 µm deep) after 1 h of incubation. Z-stack images were obtained by confocal microscopy using 60x objective. The video was realized by using ImageJ software -3D viewer plugin-. The paracellular location of the trypomastigote is shown. Green: CFSE-parasites. Red: LifeAct-RFP of HeLa cells.

**S5 movie.**

The video shows the scanning of the first 20-30 µm (from the surface to the center) of a CL Brener-infected spheroid after 1 h of incubation. Z-stack images were obtained by confocal microscopy using 60x objective. The video was realized by using ImageJ software. White arrow shows the paracellular located trypomastigote. Green: CFSE-trypomastigotes. Gray: HeLa cells.

**S6 movie**

3D reconstruction of a single intracellular CL Brener trypomastigote located inside the spheroid (20-30 µm deep) after 1 h of incubation. Z-stack images were obtained by confocal microscopy using 60x objective. The video was realized by using ImageJ software -3D viewer plugin-. Green: CFSE-trypomastigotes. Red: LifeAct-RFP of HeLa cells.

**S7 and S8 movies**

3D reconstruction of a HeLaR2 spheroid infected with CFSE-Y trypomastigotes (video 7) or CFSE-RA trypomastigotes (video 8). Z-stack images were obtained by confocal microscopy using 10x objective. The video was realized by using ImageJ software -3D viewer plugin-. A half of an infected spheroids rotating on the Y axis shown. Green: CFSE-parasites. Red: LifeAct-RFP of HeLa cells.

## References

1. OPS OMS | Enfermedad de Chagas. 2015. Available from: http://www.paho.org/hq/index.php?option=com_topics&view=article&id=10&Itemid=40743&lang=es

2. Andrade LO, Andrews NW. Opinion: The Trypanosoma cruzi–host-cell interplay: location, invasion, retention. Nat Rev Microbiol. 2005; 3(10):819– 23.

3. Rassi A, Rassi A, Marin-Neto JA, al. et, al. et, Guillen I de. Chagas disease. Lancet. 2010; 375(9723):1388–402.

4. Lewis MD, Francisco AF, Taylor MC, Jayawardhana S, Kelly JM. Host and parasite genetics shape a link between Trypanosoma cruzi infection dynamics and chronic cardiomyopathy. Cell Microbiol 2016; 18(10):1429– 43.

5. Costa FC, Francisco AF, Jayawardhana S, Calderano SG, Lewis MD, Olmo F, et al. Expanding the toolbox for Trypanosoma cruzi: A parasite line incorporating a bioluminescence-fluorescence dual reporter and streamlined CRISPR/Cas9 functionality for rapid in vivo localisation and phenotyping. PLoS Negl Trop Dis. 2018; 12(4):e0006388.

6. Prado CM, Jelicks LA, Weiss LM, Factor SM, Tanowitz HB, Rossi MA. The vasculature in chagas disease. Adv Parasitol. 2011; 76:83–99.

7. Coates BM, Sullivan DP, Makanji MY, Du NY, Olson CL, Muller WA, et al. Endothelial Transmigration by Trypanosoma cruzi. PLoS One. 2013; 8(12):e81187.

8. Lewis MD, Fortes Francisco A, Taylor MC, Burrell-Saward H, McLatchie AP, Miles MA, et al. Bioluminescence imaging of chronic Trypanosoma cruzi infections reveals tissue-specific parasite dynamics and heart disease in the absence of locally persistent infection. Cell Microbiol. 2014; 16(9):1285–300.

9. Santi-Rocca J, Gironès N, Fresno M. Multi-Parametric Evaluation of Trypanosoma cruzi Infection Outcome in Animal Models. In: Methods in molecular biology (Clifton, NJ). 2019. p. 187–202.

10. Shamir ER, Ewald AJ. Three-dimensional organotypic culture: experimental models of mammalian biology and disease. Nat Rev Mol Cell Biol. 2014; 15(10):647–64.

11. Fennema E, Rivron N, Rouwkema J, van Blitterswijk C, de Boer J. Spheroid culture as a tool for creating 3D complex tissues. Trends Biotechnol. 2013; 31(2):108–15.

12. Zanoni M, Piccinini F, Arienti C, Zamagni A, Santi S, Polico R, et al. 3D tumor spheroid models for in vitro therapeutic screening: a systematic approach to enhance the biological relevance of data obtained. Sci Rep. 2016; 6(1):19103.

13. Baruzzi A, Remelli S, Lorenzetto E, Sega M, Chignola R, Berton G. Sos1 Regulates Macrophage Podosome Assembly and Macrophage Invasive Capacity. J Immunol. 2015; 195(10):4900–12.

14. Röhland S, Wechselberger A, Spitzweg C, Huss R, Nelson PJ, Harz H. Quantification of in vitro mesenchymal stem cell invasion into tumor spheroids using selective plane illumination microscopy. J Biomed Opt. 2015; 20(4):040501.

15. Giannattasio A, Weil S, Kloess S, Ansari N, Stelzer EHK, Cerwenka A, et al. Cytotoxicity and infiltration of human NK cells in in vivo-like tumor spheroids. BMC Cancer. 2015; 15(1):351.

16. Garzoni LR, Adesse D, Soares MJ, Rossi MID, Borojevic R, Meirelles M de NL de. Fibrosis and Hypertrophy Induced by Trypanosoma cruzi in a Three-Dimensional Cardiomyocyte-Culture System. J Infect Dis. 2008; 197(6):906–15.

17. M. Ferrão P, M. Nisimura L, C. Moreira O, G. Land M, Pereira MC, de Mendonça-Lima L, et al. Inhibition of TGF-β pathway reverts extracellular matrix remodeling in T. cruzi -infected cardiac spheroids. Exp Cell Res. 2018; 362(2):260–7.

18. Cosentino RO, Agüero F. A Simple Strain Typing Assay for Trypanosoma cruzi: Discrimination of Major Evolutionary Lineages from a Single Amplification Product. PLoS Negl Trop Dis. 2012; 6(7):e1777.

19. Bernabó G, Levy G, Ziliani M, Caeiro LD, Sánchez DO, Tekiel V. TcTASV-C, a Protein Family in Trypanosoma cruzi that Is Predominantly Trypomastigote-Stage Specific and Secreted to the Medium. PLoS One. 2013; 8(7):e71192.

20. Rodríguez ME, Rizzi M, Caeiro L, Masip Y, Sánchez DO, Tekiel V. Transmigration of Trypanosoma cruzi Trypomastigotes through 3D Spheroids Mimicking Host Tissues. In: Methods in molecular biology (Clifton, NJ). 2019. p. 165–77.

21. Caeiro LD, Alba-Soto CD, Rizzi M, Solana ME, Rodriguez G, Chidichimo AM, et al. The protein family TcTASV-C is a novel Trypanosoma cruzi virulence factor secreted in extracellular vesicles by trypomastigotes and highly expressed in bloodstream forms. PLoS Negl Trop Dis. 2018; 12(5):e0006475.

22. Gerber PP, Cabrini M, Jancic C, Paoletti L, Banchio C, von Bilderling C, et al. Rab27a controls HIV-1 assembly by regulating plasma membrane levels of phosphatidylinositol 4,5-bisphosphate. J Cell Biol. 2015; 209(3):435–52.

23. Carlsson J, Yuhas JM. Liquid-overlay culture of cellular spheroids. Recent Results Cancer Res. 1984; 95:1–23.

24. Miller SA, Dykes DD, Polesky HF. A simple salting out procedure for extracting DNA from human nucleated cells. Nucleic Acids Res. 1988; 16(3):1215.

25. Campo VA. Comparative effects of histone deacetylases inhibitors and resveratrol on Trypanosoma cruzi replication, differentiation, infectivity and gene expression. Int J Parasitol Drugs drug Resist. 2017; 7(1):23–33.

26. Ramakers C, Ruijter JM, Deprez RHL, Moorman AF. Assumption-free analysis of quantitative real-time polymerase chain reaction (PCR) data. Neurosci Lett. 2003; 339(1):62–6.

27. Schneider CA, Rasband WS, Eliceiri KW. NIH Image to ImageJ: 25 years of image analysis. Nat Methods. 2012; 9(7):671–5.

28. Zingales B, Miles MA, Campbell DA, Tibayrenc M, Macedo AM, Teixeira MMG, et al. The revised Trypanosoma cruzi subspecific nomenclature: Rationale, epidemiological relevance and research applications. Infect Genet Evol. 2012; 12(2):240–53.

29. Marinho CRF, Nuñez-Apaza LN, Bortoluci KR, Bombeiro AL, Bucci DZ, Grisotto MG, et al. Infection by the Sylvio X10/4 clone of Trypanosoma cruzi: relevance of a low-virulence model of Chagas’ disease. Microbes Infect. 2009; 11(13):1037–45.

30. Marinho CRF, Nuñez-Apaza LN, Martins-Santos R, Bastos KRB, Bombeiro AL, Bucci DZ, et al. IFN-gamma, But Not Nitric Oxide or Specific IgG, is Essential for the In vivo Control of Low-virulence Sylvio X10/4 Trypanosoma cruzi Parasites. Scand J Immunol. 2007; 66(2–3):297–308.

31. Belew AT, Junqueira C, Rodrigues-Luiz GF, Valente BM, Oliveira AER, Polidoro RB, et al. Comparative transcriptome profiling of virulent and non-virulent Trypanosoma cruzi underlines the role of surface proteins during infection. PLoS Pathog. 2017; 13(12):e1006767.

32. Volta BJ, Bustos PL, Cardoni RL, De Rissio AM, Laucella SA, Bua J. Serum Cytokines as Biomarkers of Early Trypanosoma cruzi infection by Congenital Exposure. J Immunol. 2016; 196(11):4596–602.

33. Santi-Rocca J, Gironès N, Fresno M. Multi-Parametric Evaluation of Trypanosoma cruzi Infection Outcome in Animal Models. 2019. p. 187–202.

34. Lewis MD, Francisco AF, Taylor MC, Kelly JM. A New Experimental Model for Assessing Drug Efficacy against Trypanosoma cruzi Infection Based on Highly Sensitive In Vivo Imaging. J Biomol Screen. 2015; 20(1):36–43.

35. Harker KS, Ueno N, Lodoen MB. Toxoplasma gondii dissemination: a parasite’s journey through the infected host. Parasite Immunol. 2015; 37(3):141–9.

36. Kumar H, Tolia NH. Getting in: The structural biology of malaria invasion. PLOS Pathog. 2019; 15(9):e1007943.

37. Drewry LL, Sibley LD. The hitchhiker’s guide to parasite dissemination. Cell Microbiol. 2019; e13070.

38. Lambert H, Barragan A. Modelling parasite dissemination: host cell subversion and immune evasion by Toxoplasma gondii. Cell Microbiol. 2010 Mar; 12(3):292–300.

39. Mulenga C, Mhlanga JD, Kristensson K, Robertson B. Trypanosoma brucei brucei crosses the blood-brain barrier while tight junction proteins are preserved in a rat chronic disease model. Neuropathol Appl Neurobiol. 2001; 27(1):77–85.

40. Grab DJ, Nikolskaia O, Kim Y V., Lonsdale-Eccles JD, Ito S, Hara T, et al. African trypanosome interactions with an in vitro model of the human blood-brain barrier. J Parasitol. 2004; 90(5):970–9.

41. Nikolskaia O V., de A Lima APC, Kim Y V, Lonsdale-Eccles JD, Fukuma T, Scharfstein J, et al. Blood-brain barrier traversal by African trypanosomes requires calcium signaling induced by parasite cysteine protease. J Clin Invest. 2006; 116(10):2739–47.

42. Weight CM, Jones EJ, Horn N, Wellner N, Carding SR. Elucidating pathways of Toxoplasma gondii invasion in the gastrointestinal tract: involvement of the tight junction protein occludin. Microbes Infect. 2015; 17(10):698–709.

43. Barragan A, Brossier F, Sibley LD. Transepithelial migration of Toxoplasma gondii involves an interaction of intercellular adhesion molecule 1 (ICAM-1) with the parasite adhesin MIC2. Cell Microbiol. 2005; 7(4):561–8.

44. Huynh M-H, Carruthers VB. Toxoplasma MIC2 Is a Major Determinant of Invasion and Virulence. PLoS Pathog. 2006; 2(8):e84.

45. Yang ASP, O’Neill MT, Jennison C, Lopaticki S, Allison CC, Armistead JS, et al. Cell Traversal Activity Is Important for Plasmodium falciparum Liver Infection in Humanized Mice. Cell Rep. 2017; 18(13):3105–16.

46. Barragan A, Sibley LD. Transepithelial migration of Toxoplasma gondii is linked to parasite motility and virulence. J Exp Med. 2002 Jun 17;195(12):1625–33.

47. Howe DK, Sibley LD. Toxoplasma gondii Comprises Three Clonal Lineages: Correlation of Parasite Genotype with Human Disease. J Infect Dis. 1995; 172(6):1561–6.

48. Fuentes I, Rubio JM, Ramirez C, Alvar J. Genotypic Characterization of Toxoplasma gondii Strains Associated with Human Toxoplasmosis in Spain: Direct Analysis from Clinical Samples. J Clin Microbiol. 2001; 39(4):1566–70.

49. Dóró É, Jacobs SH, Hammond FR, Schipper H, Pieters RP, Carrington M, et al. Visualizing trypanosomes in a vertebrate host reveals novel swimming behaviours, adaptations and attachment mechanisms. Elife. 2019; 8. e48388.

50. Castillo C, Carrillo I, Libisch G, Juiz N, Schijman A, Robello C, et al. Host-parasite interaction: changes in human placental gene expression induced by Trypanosoma cruzi. Parasit Vectors. 2018;11(1):479.

51. Juiz NA, Torrejón I, Burgos M, Torres AMF, Duffy T, Cayo NM, et al. Alterations in Placental Gene Expression of Pregnant Women with Chronic Chagas Disease. Am J Pathol. 2018; 188(6):1345–53.

52. Bustos PL, Milduberger N, Volta BJ, Perrone AE, Laucella SA, Bua J. Trypanosoma cruzi Infection at the Maternal-Fetal Interface: Implications of Parasite Load in the Congenital Transmission and Challenges in the Diagnosis of Infected Newborns. Front Microbiol. 2019; 10:1250.

53. Juiz NA, Solana ME, Acevedo GR, Benatar AF, Ramirez JC, da Costa PA, et al. Trypanosoma cruzi prod1. Juiz NA, Solana ME, Acevedo GR, Benatar AF, Ramirez JC, da Costa PA, et al. Different genotypes of Trypanosoma cruzi produce distinctive placental environment genetic response in chronic experimental infecti. PLoS Negl Trop Dis. 2017; 11(3):e0005436.

54. Stuelten CH, Parent CA, Montell DJ. Cell motility in cancer invasion and metastasis: insights from simple model organisms. Nat Rev Cancer. 2018; 18(5):296–312.

55. Wolf K, Friedl P. Extracellular matrix determinants of proteolytic and non-proteolytic cell migration. Trends Cell Biol. 2011; 21(12):736–44.

